# Genomic investigation of a suspected multi-drug resistant *Klebsiella pneumoniae* outbreak in a neonatal care unit in sub-Saharan Africa

**DOI:** 10.1101/2020.08.06.236117

**Authors:** Jennifer Cornick, Patrick Musicha, Chikondi Peno, Ezgi Saeger, Pui-ying Iroh Toh, Aisleen Bennett, Neil Kennedy, Nicholas Feasey, Eva Heinz, Amy K. Cain

**Author notes:** Corresponding authors. Emails: Amy K. Cain and Jennifer Cornick.

## Abstract

A suspected outbreak of multi-drug resistant (MDR) *Klebsiella pneumoniae* in a Malawian neonatal unit was investigated using whole-genome sequencing. Strain-types, virulence and resistance genes of *K. pneumoniae* isolated from patients from the hospital over a four-year period were identified. A MDR ST340 clone was implicated as the likely outbreak cause.

---

*Klebsiella pneumoniae* is an opportunistic pathogen, responsible for an increasing burden of hospital-acquired infections globally. It has the ability to readily acquire antimicrobial resistance mechanisms, with sequence types (STs), ST258, ST11, ST14/15 and ST405, reported to cause a large proportion of cephalosporin- and carbapenem-resistant infections with high mortality [1]. Critically, sepsis caused by multi-drug resistant (MDR) *K. pneumoniae* is a growing challenge in neonatal care units; a recent systematic review of extended spectrum beta-lactamase (ESBL) producing Enterobacteriaceae in neonatal care units reported that ESBL *K. pneumoniae* is the pathogen most frequently responsible for outbreaks in these settings, and is associated with a mortality rate of 31% [2]. There are multiple reports detailing the prevalence and genetic epidemiology of MDR *K. pneumoniae* causing hospital outbreaks in high-income settings [3]. However, reports from low- and middle-income settings, where such data are of immense value in implementing appropriate infection prevention control and treatment regimes, are scarce.

Chatinkha is a 70-bed neonatal special care unit at Queen Elizabeth Central Hospital (QECH), the biggest government referral hospital in Southern Malawi. Given that Chatinkha predominantly admits neonates born at QECH with complications, the majority of blood stream infections (BSI) diagnosed on this unit are considered to be hospital acquired. At QECH, all children with clinically suspected sepsis undergo a blood culture (BC) test [4]. The BC service typically identifies <2 *K. pneumoniae* BSI on Chatinkha each month, however a spike in cases was observed in February (n=7) and March (n=9) 2014. The number of cases temporarily decreased, however September to November 2014 showed a second peak in cases (Figure TA1). Despite Chatinkha having the smallest capacity of all of the paediatric wards at QECH, from February to November 2014 over 75% of all paediatric *K. pneumoniae* BSI (n=33/43) were from neonates admitted to Chatinkha. This rise was not due to an increase in the number of blood cultures taken, which remained stable (Fig 1A). Furthermore, *K. pneumoniae* BSI in Chatinkha during this period were untreatable with locally available antibiotics; most displayed identical antimicrobial resistance profiles, suggesting the possibility of a clonal outbreak.

**Figure 1.**
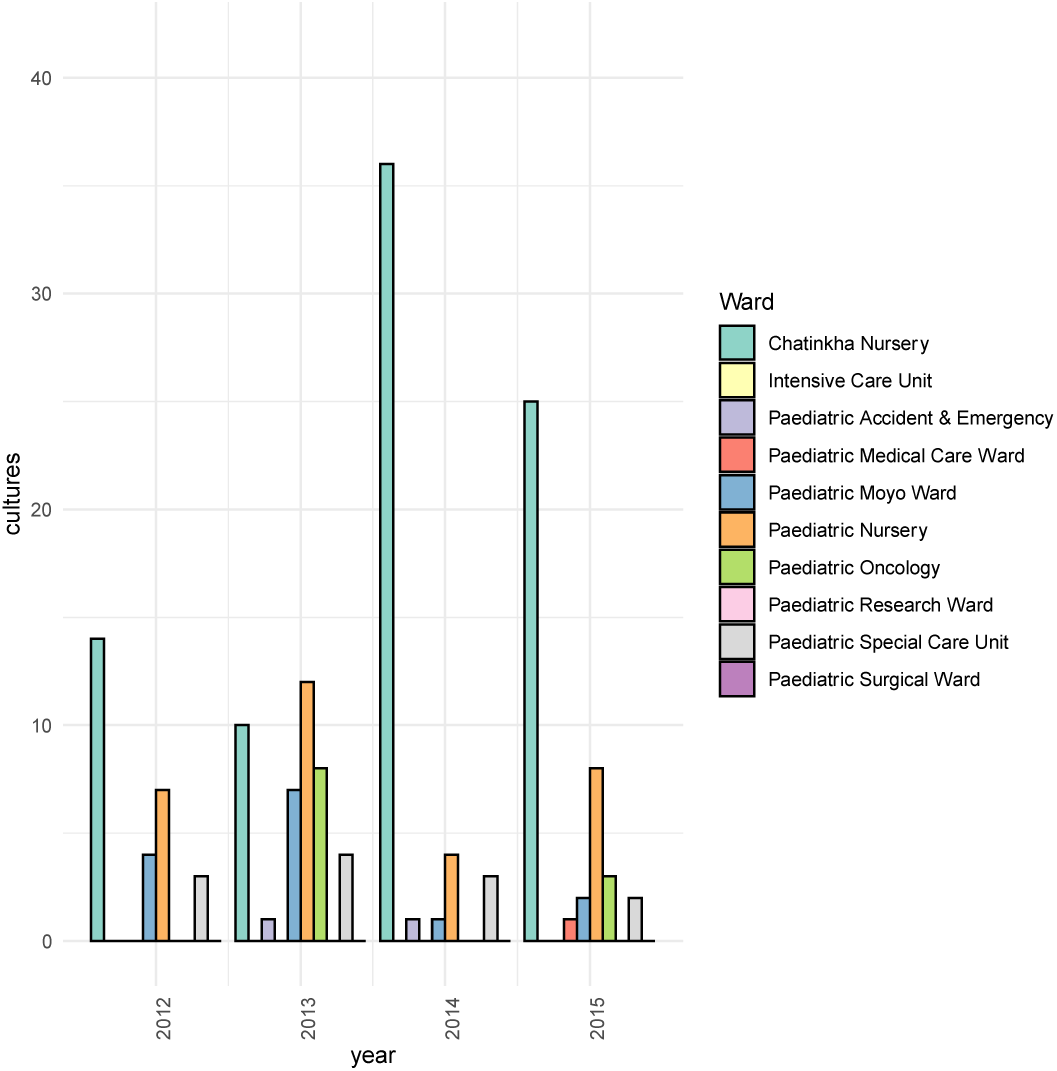
Bar chart showing the number of *Klebsiella pneumoniae* positive blood cultures for Chatinkha ward and other pedriatric wards overlaid with a scatter plot showing the total number of blood cultures (including culture negative) from the paediatric wards at Queen Elizabeth Central Hospital, Malawi from 2012 – 2015. Data on the total number of blood cultures taken was not available for 2012 and is not shown. **B. The distribution of antimicrobial resistance elements within the *Klebsiella pneumoniae* isolates from Queen Elizabeth Central Hospital**. The phylogeny is based on a core genome SNP alignment of the Malawian isolates (n=86), the branches are labeled with ST. The panel adjoining the phylogeny shows the AMR phenotype and the absence/presence of key AMR genes [*bla*_CTX-M-15_, *bla*_OXA-1_, *bla*_OXA-2_, *bla*_OXA-16_, *bla*_SHV-11_, *bla*_TEM-1_ (beta-lactams), *aac*(6’)-Ib-cr (quinolone), *str*AB (streptomycin), *sul1* (sulphonamides), *catA1, catA2* (phenicols), *tetA*(D) (tetracycline), *aadA1, aadA2, aac*(3’)-IIa (aminoglycosides), *dfrA1, dfrA2* (trimethoprim) and *mph* (macrolides)] and plasmids amongst the isolates.

Here, we performed a genomic investigation into the suspected *K. pneumoniae* outbreak, sequencing genomes from February to November 2014 (boxed in Fig TA1). All viable, archived *K. pneumoniae* BSI isolates from Chatinkha (n=62) and a subset of *K. pneumoniae* BSI isolates (n=38) from other paediatric wards isolated January 2012 to December 2015, were whole genome sequenced using the Illumina HiSeq X-Ten platform. Fourteen sequenced genomes failed initial quality control, the remaining 86 samples (Chatinkha n=56, other wards n=30) yielded on average 2.3 million reads/sample. The sequences were deposited in the European Nucleotide Archive under project numbers PRJEB19322 and PRJNA641987. Isolates were previously subjected to antibiotic susceptibility testing using the dics diffusion method and BSAC guidelines [4].

Multilocus sequence (MLST) typing was performed *in silico* as described elsewhere [5]. Resistance genes were identified using SRST2 [6], plasmids using PBRT [7], capsular type using Kaptive and virulence genes using Kleborate v0.3.0 (https://github.com/katholt/Kleborate). In order to place the study dataset within a global context, we added our genomes to a dataset from an international *K. pneumoniae* study [1]. A core gene alignment of the combined genomic dataset was generated using ROARY [8] and from this, single nucleotide variants were used to generate a phylogeny with RaxML v.7.8.6 [9]. Patient records were retrospectively analysed to establish mortality outcomes. Ethical approval for the study was awarded by the College of Medicine Research Ethics Committee (P.08/14/1614 and P.018/17/2255).

Whole genome sequencing analysis identified two lineages, ST340 and ST14, as the dominant *K. pneumoniae* STs recovered from neonates admitted to Chatinkha (Fig TA1).

## There was a discreet outbreak of MDR ST340

ST340, an ST within the CG258 complex that has been associated with MDR hospital infections worldwide [10,11], accounted for almost a third (30%, n=17/56) of all *K. pneumoniae* isolated from Chatinkha during the study period. All ST340 isolates were capsular type 15 and serotype O4 and differed from one another by ≤30 SNPs (Fig TA2), indicating that a single ST340 lineage was circulating in the ward over the entire study period. Over the suspected outbreak period, ST340 accounted for more than half of the *K. pneumoniae* BSI from Chatinkha (58%, n=14/24), strongly suggesting that dissemination of this clonal strain within Chatinkha was responsible for the peak in BSI observed from February to November 2014. The ST340 were resistant to augmentin, ampicillin, ceftriaxone, chloramphenicol, ciprofloxacin, cotrimoxazole, gentamicin and susceptible to amikacin, an antibiotic not local availabile. Consistent with this, all ST340 isolates harboured multiple antibiotic resistance genes, conserved between all ST340 isolates, namely *bla*_CTX-M-15_, *bla*_OXA-1_, *bla*_OXA-2_, *bla*_OXA-16_, *bla*_SHV-11_, *bla*_TEM-1_ (beta-lactams), *aac*(6’)-Ib-cr (quinolone), *str*AB (streptomycin), *sul1* (sulphonamides), *catA1, catA2* (phenicols), *tetA*(D) (tetracycline), *aadA1, aadA2, aac*(3’)-IIa (aminoglycosides), *dfrA1, dfrA2* (trimethoprim) and *mph* (macrolides) and a number of plasmids (see Fig 1B). Of the ST340 BSI cases reported in neonates on Chatinkha, outcome data was available for 13, of which four died (case fatility rate, 31%).

Prior to the outbreak, ST340 BSI had only been isolated from Chatinkha on two previous occasions (May and November 2013) (Fig TA1). These isolates differed from the first two ST340 cases reported in February 2014 by <5 SNPs (Fig TA2), confirming that closely related strains were circulating in Chatinkha at least six-months prior to the outbreak. Previous to this, ST340 was isolated on a single occasion from two other wards, the earliest in February 2013. These two ST340s isolates differed from the presumed outbreak ST340 precursor isolate by <7 SNPs. Whilst we cannot confirm the exact date at which ST340 was seeded into QECH, this indicates that ST340 was circulating in the hospital at least a year prior to the outbreak. Following the outbreak, ST340 was identified twice more on Chatinkha. A single case in April 2015 and again a single case in October 2015. October 2015 saw six cases of *K. pneumoniae* BSI identified on Chatinkha, hinting at the start of another outbreak, however these six cases were caused by five different STs. The fact ST340 continued to circulate in the ward post November 2014 but did not to contribute to a further peak in cases, suggests that the success of this clone during the outbreak period was not driven by genomic factors alone.

## ST14 did not contribute to the outbreak

Despite being the second most commonly isolated ST from Chatinkha (27%, n=15/56) during the study period, only a single ST14 case was reported during the outbreak period, in March 2014. ST14 showed a greater level of variation in their resistance profiles relative to ST340, and isolates were predominantly susceptible to Chloramphenicol and Ciprofloxacin as well as Amikacin (Fig TA2).

## ST372 related to a peak in cases

ST372 (capsular type: 43, serotype: O2V1) also appeared to contribute to the peak in *K. pneumoniae* BSI over the outbreak period as ST372 was responsible for 20.8% (n=5/24) of cases. ST372 was first observed in February 2014 in Chatinkha and caused four BSI in one month and the remaining five *K. pneumoniae* BSIs reported during the outbreak period belonged to five different STs.

## The outbreak strains in a global context

A core phylogeny was constructed to place isolates into a global context using a sequencing dataset of diverse *K. pneumoniae* isolates [13] (Fig TA3). Although no ST340 isolates were present in this dataset, the Malawian ST340 were most closely related to ST258 isolates from the USA. Interestingly, ST14 strains from Malawi were clonally related (<100 SNPs) to international isolates with >99.9% identity to isolates from Australia and the Netherlands, indicating a global spread.

## Conclusion

We studied *K. pneumoniae* BSI isolated from neonates admitted to neonatal unit over a four-year period, encompassing a suspected outbreak in 2014, as well as a representative subset of *K. pneumoniae* BSI reported from other paediatric wards within the same hospital.

We show that ST340, and to a lesser extent ST372, caused an increase in BSI reported on the neonatal ward in 2014. ST340 was observed in other wards prior to the outbreak, suggesting that it was circulating in the hospital prior to the outbreak. ST372 was not identified in other wards prior to the outbreak, however this may be due to the limitations of our sampling. In addition to the outbreak lineages we observed a large cluster of ST14 isolates, which intermittently contributed to no more than two BSI cases per month on Chatinkha. All other ST identified in Chatinkha over the sampling period were only observed in sporadic single isolate clusters. From the sequencing analysis alone, it is unclear what exact factors allowed the specific lineages to persist or successfully transmit within Chatinkha beyond antibiotic selection and this is under further investigation. Further, continual monitoring of the hospital environment, especially for the re-emergence of MDR ST340, is urgently needed in order to implement measures to prevent the persistence and/or spread of these locally untreatable lineages.

## ABBREVIATIONS

(BSI): Blood stream infection;
(ESBL): Extended spectrum beta-lactamase;
(MDR): Multi-drug resistance;
(MLST): Multi locus sequence type;
(ST): Sequence type;
(QECH): Queen Elizabeth Central Hospital.

## TECHNICAL APPENDIX

### (**DETAILED METHODS**)

The blood culture and CSF data presented here were collected at QECH, which serves both as district hospital for Blantyre and tertiary hospital for the surrounding districts. The hospital admits approximately 10,000 adult (aged ≥16 years) and 50,000 pediatric (aged< 16 years) medical patients/year. Blood culture collection, processing and antimicrobial susceptibility testing methods for these isolates have been previously described [4].

**TA1.**
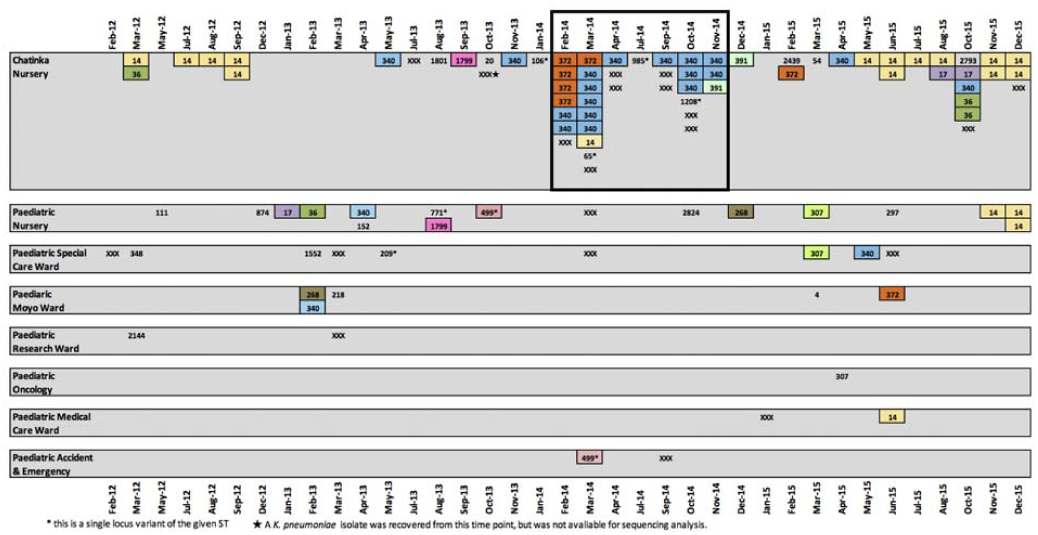
Schematic showing the location, sequence type (ST) and date of isolation of the *K. pneumoniae* isolates that were subjected to whole genome sequencing (n=86). STs that were identified more than once in the dataset are highlighted with a coloured box reflecting the STs. Singleton STs are not highlighted. Non-viable isolates (n=4) and those that failed sequencing (n=14) but were earlier confirmed as *K. pneumoniae* by the diagnostic laboratory are marked on the schematic as ‘XXX’ in order to give complete picture of the number of BSI *K. pneumoniae* cases reported from Chatinkha each month during the study period. Months where no cases were reported are omitted from the schematic. The suspected outbreak period is highlighted with a black outline.

**Figure TA2.**
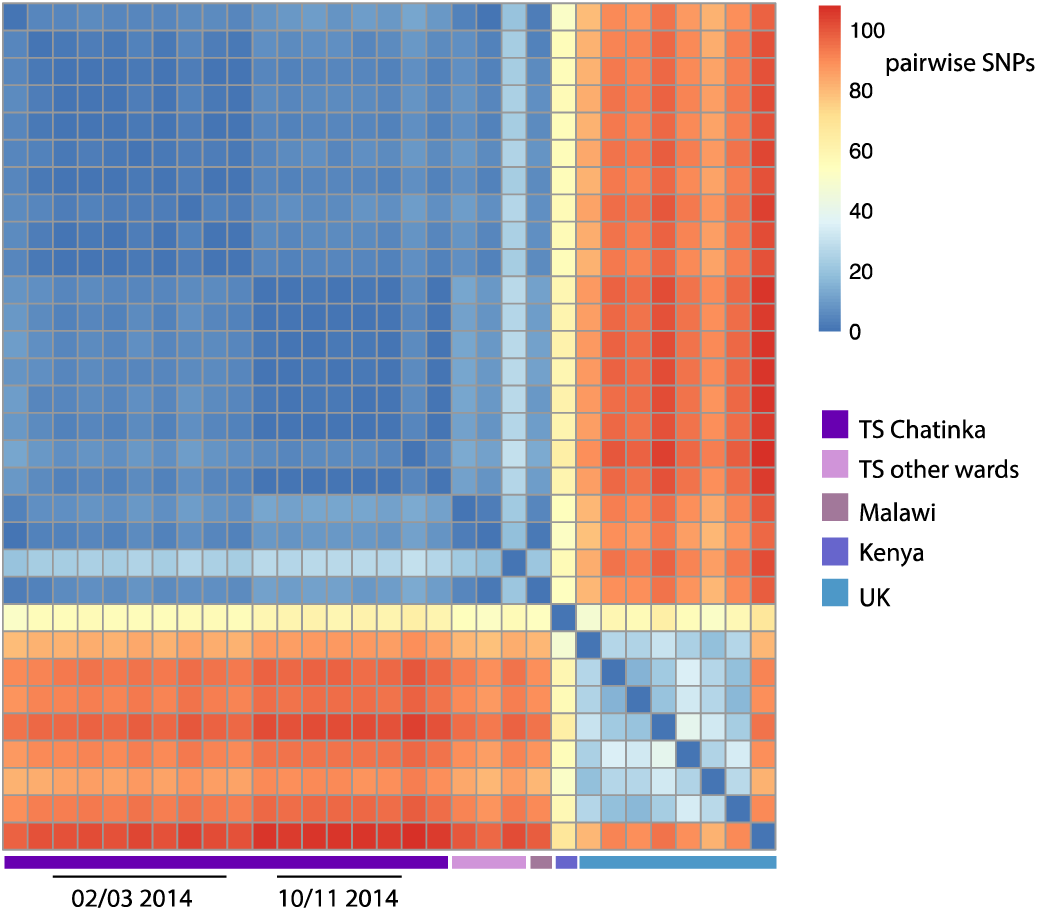
Heatmap showing the number of SNP differences between the *K. pneumoniae* ST340 isolates sequenced as part of the outbreak investigation. To bring the relatedness of our samples into context with other ST340 isolates, the analysis includes a community acquired Malawian ST340 sequenced as part of a previous study at QECH [4], a single ST340 isolate from a study in Kenya [14], and eight ST340 genomes from the UK [14].

**Figure TA3.**
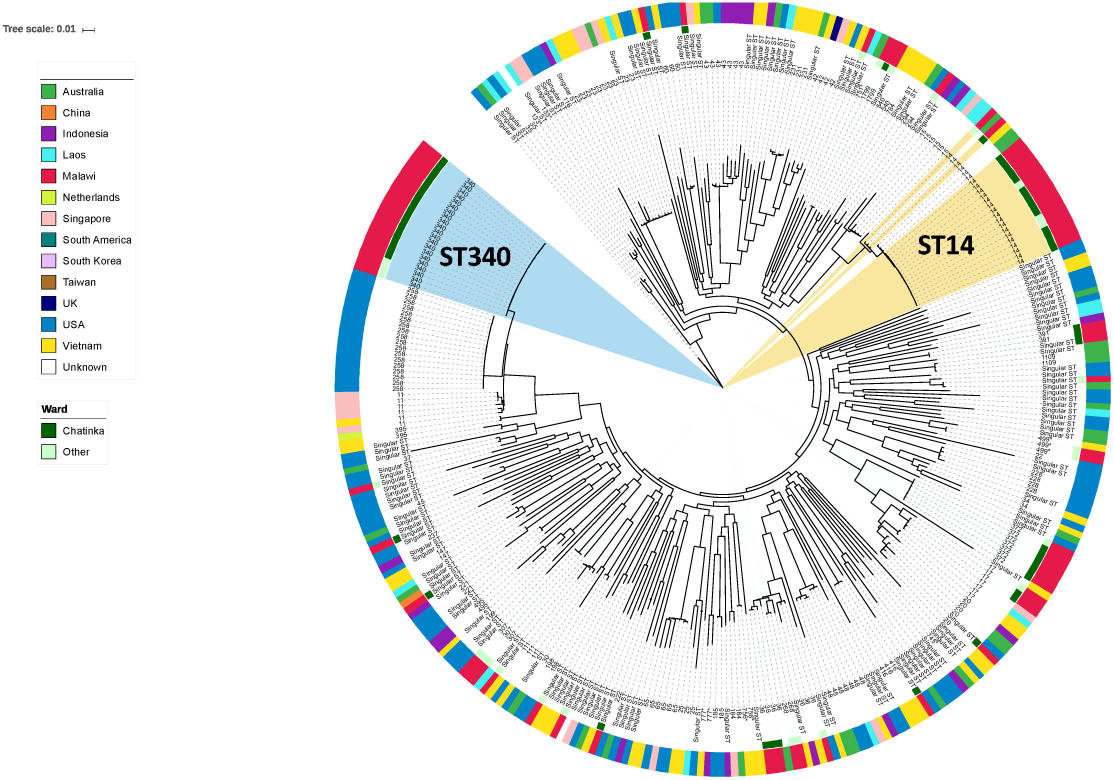
Population structure of Klebsiella pneumoniae. A core genome phylogeny of the Malawian KP-I isolates (n=81) in the context of a previously published global dataset. Branch labels are annotated with ST. The inner colored circle indicates if the Malawian isolates were recovered from the Chatinkha neonatal care unit, the outer ring indicates the country of isolation.

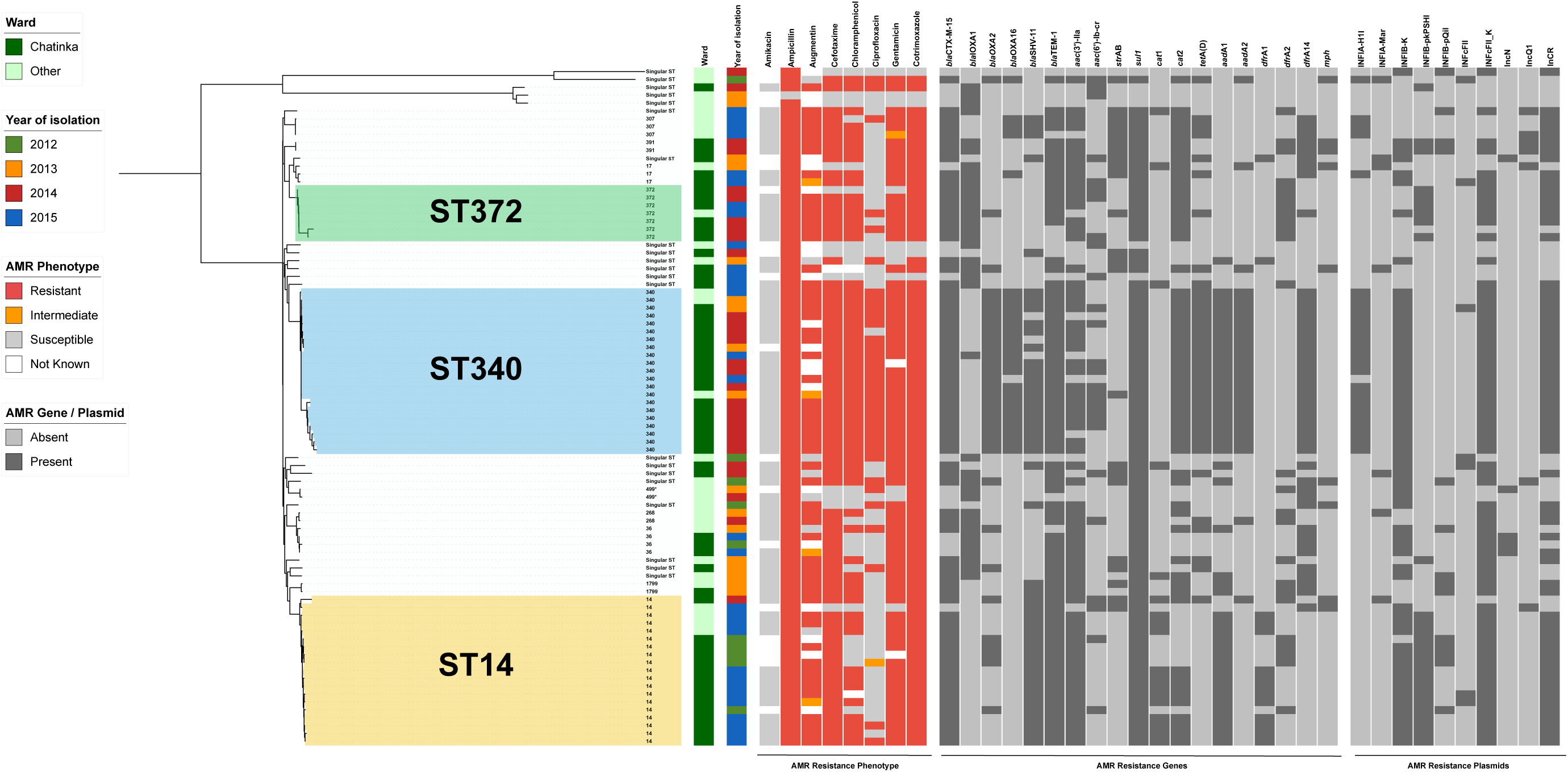

